# Hippocampal Development in a Rat Model of Perigestational Opioid Exposure

**DOI:** 10.64898/2026.03.29.715159

**Authors:** Meghan E. Vogt, Jade Kang, Anne Z. Murphy

**Affiliations:** Neuroscience Institute, Georgia State University, 100 Piedmont Ave Se., Atlanta, GA 30303

**Author notes:** **Corresponding Author:** Anne Z. Murphy, 100 Piedmont Ave., Atlanta, GA, 30303. The authors declare no competing financial interests.

## Abstract

Nearly one third of women of reproductive age in the United States are prescribed opioids annually; 14% of women fill an opioid prescription during pregnancy, and one in five report misuse. Opioid use during pregnancy has given rise to an increasing population of infants born with gestational opioid exposure. Although substantial clinical work has focused on treating these infants as they experience opioid withdrawal symptoms at the time of birth, notably few studies have examined the effects of gestational opioid exposure on brain development and long-term cognitive function. During typical brain development, endogenous opioids and their receptors are highly expressed by neural progenitor cells, neurons, and glia where they modulate cell proliferation, differentiation, and maturation. Thus, any disruption to the endogenous opioid system during the critical period of brain development may have lasting consequences on brain cell populations and the behaviors they influence. Indeed, opioid-exposed infants have smaller brains than age-matched peers and show significant neurodevelopmental impairment; they also have higher rates of learning disability at school age. To investigate how exposure to exogenous opioids during brain development affects neural maturation in the hippocampus, a brain region critical for learning and memory, our lab has developed a clinically relevant perigestational morphine exposure rat model. The current study reports that perigestational exposure to morphine delays postnatal hippocampal neuronal maturation, alters astrocyte and oligodendrocyte proliferation, and alters expression of brain-derived neurotrophic factor (BDNF), a protein crucial for healthy brain growth. Furthermore, we show that environmental enrichment rescues BDNF deficits, offering evidence for the effectiveness of non-invasive, non-pharmacological intervention for developmental consequences of perigestational opioid exposure.

## Introduction

Over the past two decades, opioid use during pregnancy has become increasingly common in the United States; indeed, nearly one third of reproductive-age women are prescribed opioids, and an estimated 14-22% fill an opioid prescription during pregnancy (Ailes et al., 2015; Ko et al., 2020). As opioids readily cross the placenta, gestational use impacts the developing fetus as well as the mother.

Neural progenitor cells, neurons, and glia express high levels of endogenous opioids and their receptors across development (Hauser & Knapp, 2018; Kornblum et al., 1987; Leslie & Loughlin, 1993; Zhu et al., 1998; Leslie et al., 1998; Sargeant et al., 2007). Such high expression allows the endogenous opioid system to modulate many developmental processes, including neural proliferation, differentiation, and maturation. Interestingly, even brain regions that do not express opioid receptors in adulthood show expression during development, highlighting the key role endogenous opioid signaling plays in organizing the early nervous system (Spodnick et al., 2025; Reznikov et al., 1999; Stiene-Martin et al., 2001). Thus, the developing brain is uniquely susceptible to disruption by exogenous opioids introduced by maternal drug use.

Clinically, infants with gestational opioid exposure have significantly smaller total brain volumes and region-specific abnormalities that persist after accounting for maternal factors, such as tobacco use and education (Radhakrishnan et al., 2021a; Wu et al., 2025; Madurai & Jantzie, 2025). Infants with gestational opioid exposure also display general white matter abnormalities and punctate white matter lesions that have been associated with motor deficits, delayed speech, and delay in meeting developmental milestones (Merhar et al., 2019; Nguyen et al., 2019; Merhar et al., 2025). At school age, these children show learning deficits, repeating grades at a higher rate than non-exposed controls and often requiring speech therapy to improve verbal impairments (Hunt et al., 2008; Fill et al., 2018; Maguire et al., 2016; Sundelin Wahlsten & Sarman, 2013; Wilson et al., 1979). Furthermore, adolescents with prenatal opioid exposure demonstrate impaired performance and altered prefrontal cortex activation when completing the most demanding conditions of a working memory-selective attention task (Sirnes et al., 2018). Together, these morphological and cognitive studies highlight the developmental consequences of children exposed to opioids *in utero,* however, the literature lacks a clear mechanistic understanding of how opioid exposure during development disrupts neurodevelopment.

To address this gap, our lab has developed an opioid exposure model that recapitulates a clinical timeline in which adolescent female rats receive morphine through daily, intermittent administration that begins before pregnancy, continues through gestation, and is tapered postnatally (Substance Abuse and Mental Health Services Administration, 2014).

We have reported reduced juvenile play in opioid-exposed offspring (Harder et al., 2023) and reductions in alcohol and sucrose preference (Searles et al., 2025). Relevant to the present study, we showed that perigestational morphine exposure impairs hippocampus-dependent spatial learning. Specifically, male and female adolescent rats who were exposed to morphine relied significantly less on the hippocampal-dependent spatial strategy of the Barnes maze. However, the mechanism by which opioid exposure during development could impact hippocampal-dependent performance remains unknown. Thus, the present study tested the hypothesis that perigestational morphine exposure impairs hippocampal development; we investigated neuronal and glial proliferation and maturation as well as BDNF expression. As previous studies have reported significant increases in such developmental measures following environmental enrichment, additional experiments assessed the ability of environmental enrichment to rescue the observed deficits following perigestational morphine exposure.

## Methods

### Subjects

Female Sprague Dawley rats (age P60; Charles River Laboratories) were bred with sexually experienced males for offspring generation. Offspring were weaned at P21 into Optirat GenII individually ventilated cages (Animal Care Systems) with corncob bedding (Bed-o’Cobs; The Andersons). Rats were housed in same-sex, same-treatment groups of two to four on a 12/12-hour light/dark cycle (lights off at 8:00 AM). Rodent chow (Laboratory Rodent Diet 5001 or 5015 for breeding pairs; Lab Diet) and water were provided *ad libitum* in the home cage. Experiments were approved by the Institutional Animal Care and Use Committee at Georgia State University and performed in compliance with the National Institutes of Health Guide for the Care and Use of Laboratory Animals. All efforts were made to reduce the number of rats used and minimize pain and suffering.

### iPrecio Pump Implantation Surgery & Perigestational Morphine Exposure

Microinfusion pumps (iPrecio SMP-200; Alzet) were surgically implanted into female rats at postnatal day 60 (P60) as previously described (Vogt & Murphy, 2025). Before implantation, pumps were programmed to deliver 30µL of solution across one hour three times per day (12:00 PM, 8:00 PM, 4:00 AM); the flowrate during non-infusion was 0.1µL/hour. Rats were randomly assigned to either perigestational morphine exposure (MOR) or control (VEH). Morphine administration was initiated one-week post-surgery. One week following morphine initiation, females were paired with sexually experienced males for two weeks to increase chance of pregnancy. Morphine (or saline) continued throughout gestation. Rats were initially administered 10mg morphine per kg body weight per day, with doses increasing weekly by 2 mg/kg/day until 16 mg/kg/day was reached. At approximately embryonic day 18 (E18), infusions decreased to twice-a-day to minimize early-life pup mortality. Post-parturition, dams continued to receive decreasing doses of morphine until P7 when administration ceased, and morphine was replaced with sterile saline. This protocol closely mirrors the clinical profile of a late adolescent female who is regularly using an opioid, becomes pregnant, and continues using throughout pregnancy and following birth.

### Immunohistochemistry

On postnatal days 7, 14, and 30, brains of morphine- (MOR) or saline-exposed (VEH) offspring were rapidly extracted following decapitation and stored in 4% paraformaldehyde at 4°C for 24 hours, then transferred into 30% sucrose until sectioning. Fixed tissue was sectioned coronally at 40µm with a Leica SM2010R microtome; sections were stored at −20°C in cryoprotectant solution until immunohistochemical experiments (Watson et al., 1986).

For chromogen staining to assess neuronal maturation, free-floating sections were rinsed in potassium phosphate-buffered saline (KPBS), incubated in 3% hydrogen peroxide for 30 minutes, then rinsed again before incubation in mouse anti-NeuN primary antibody (1:100,000; MAB377; Millipore) for 48 hours at 4°C. Following thorough rinsing, tissue was incubated in biotinylated donkey anti-mouse IgG (1:600; 715-065-151; Jackson ImmunoResearch) at room temperature for 1 hour, rinsed, and then secondary antibody signal was amplified with avidin-biotin solution (AB; 45μL each per 10mL; PK-6100; Vector Laboratories) for 1 hour. After AB incubation, tissue was rinsed with KPBS and sodium acetate (0.175M) before immunoreactivity was visualized using nickel sulfate 3,3′-diaminobenzidine solution (DAB; 2mg/10ml) activated with 0.08% hydrogen peroxide in sodium acetate buffer. After 15 minutes of DAB incubation, tissue was rinsed with sodium acetate and KPBS, mounted onto gelatin-subbed slides, and dehydrated with increasing concentrations of ethanol before cover-slipping.

For fluorescent staining, free-floating sections were rinsed in KPBS, then antigen retrieval was performed by incubation in EDTA buffer at 90°C for three minutes. After thorough rinsing, tissue was blocked with 5% normal donkey serum (017-000-121; Jackson ImmunoResearch) for 1 hour at room temperature before antibody incubations. Tissue was mounted on gelatin-subbed slides and cover-slipped with ProLong Diamond Antifade Mountant (Thermo Fisher Scientific).

To identify newly proliferated, immature neurons, tissue was incubated in mouse anti-Tubβ3 (1:2000; sc-80016; Santa Cruz Biotechnology) for 1 hour at room temperature then overnight at 4°C. Tissue was rinsed and then incubated in donkey anti-mouse Alexa Fluor 555 secondary antibody (1:500; A31570; Thermo Fisher Scientific) and 0.02% DAPI (62247; Thermo Fisher Scientific) for 2 hours at room temperature.

To label newly proliferated cells, tissue was incubated in rabbit anti-Ki67 (1:500; AB9260; Abcam) for 1 hour at room temperature then overnight at 4°C. Tissue was rinsed and then incubated in donkey anti-rabbit Alexa Fluor 488 secondary antibody (1:300; A21206; Thermo Fisher Scientific) for 2 hours at room temperature followed by thorough rinsing. To label astrocytes, tissue was incubated in rabbit anti-S100β (1:8000; AB52642; Abcam) for 1 hour at room temperature and then overnight at 4°C. Tissue was rinsed and then incubated in donkey anti-rabbit Alexa Fluor 555 secondary antibody (1:500; A31572; Thermo Fisher Scientific) and 0.02% DAPI for 2 hours at room temperature. In some assays, rabbit anti-Olig2 (1:1000; AB109186; Abcam) was used instead of Ki67 & S100β to label oligodendrocyte precursor cells and early oligodendrocytes.

### Imaging & Quantification

Z-stacked images were captured using a Keyence BZ-X700 microscope and maximum intensity projections were quantified using ImageJ software (Schindelin et al., 2012). Subregions of the hippocampus (HPC; 1.8-4.0mm posterior to bregma, depending on age) were bilaterally imaged at 20 – 40x magnification (2-4 images per region per subject) and background-corrected before analysis. For NeuN, images were thresholded to capture NeuN signal. Then, captured area of signal was divided by the total area of the dentate gyrus. For Tubβ3, mean fluorescent intensity within the region of interest was measured. For Olig2 and S100β, the number of positively stained cells were hand-counted and corrected for area of region quantified.

### BDNF Enzyme Linked Immunosorbent Assay

On postnatal day 14, offspring brains were rapidly extracted following decapitation and frozen on dry ice. Dorsal hippocampus (1.6-2.2mm posterior to bregma) was dissected and stored at −80°C until homogenization. Samples were homogenized with a motorized pestle in RIPA buffer (50mM Tris-HCl, 150mM NaCl, 1% Triton X-100, 0.5% sodium deoxycholate, Roche cOmplete Mini EDTA-free Protease Inhibitor Cocktail) and stored in 100mg/mL aliquots at −80°C until use in assay.

Hippocampal levels of proBDNF and mature BDNF were quantified using the Biosensis ELISA kits (BEK-2211 & 2237; Biosensis). Samples were diluted 1:20 with Sample Diluent A provided by the manufacturer; all assays were run in duplicates. Correlation coefficients were measured against known standards (r^2^ > 0.99); intra-assay CVs were below 8%.

### Environmental Enrichment

To boost sensory stimulation, litters received home cage environmental enrichment (EE) in the form of additional nesting material and shelters. Enrichment was introduced at P7 as pilot studies noted that altering the home cage at this timepoint was the least disruptive in terms of maternal stress and impact on maternal behavior. Brains were collected from MOR EE offspring at P14 and were assessed for hippocampal Olig2 immunoreactivity and BDNF expression.

### Statistical Analyses

All data met the requirements for parametric analyses (normality, homogeneity of variance) unless otherwise noted. The effects of treatment, sex, age (where applicable), and region (where applicable) were assessed using t-tests (k=2), one-way, or two-way mixed model ANOVA with an alpha level of 0.05 unless otherwise noted. When no sex difference or sex x treatment interaction was present, sexes were combined to increase power. Tukey’s or Sidak’s post hoc tests were conducted to determine significant mean differences between *a priori* specified groups. Twelve vehicle-exposed and 15 morphine-exposed litters (3 of which received enrichment) are represented in the immunohistochemistry data; 5 vehicle-exposed and 9 morphine-exposed litters (2 of which received enrichment) are represented in the BDNF data. No significant litter or cohort effects were observed when litter was run as a covariate; thus, offspring from above-mentioned litters, generated across 13 cohorts, were combined into one dataset for analysis. All image quantifications and ELISA assays were conducted blind to treatment. Graphs depict mean ± SEM unless otherwise noted. GraphPad Prism 10.5.0 was used for all statistical analyses.

## Results

### Neurogenesis

To assess the impact of perigestational morphine on postnatal hippocampal neurogenesis, brains from P7 and P14 rats were stained for beta-tublin-3 (Tubβ3), a neuron-specific cytoskeletal protein that is highly expressed shortly after proliferation with diminishing expression as the neuron matures (Kempermann 2003). We report a significant age effect of Tubβ3 signal in the hippocampus, with P7 levels averaging 56% higher than P14 [F(1,37)=10.76, p=0.0023; **Figure 1**]. No significant treatment effects were observed [F(1,37)=0.1857, p=0.6690].

**Figure 1:**
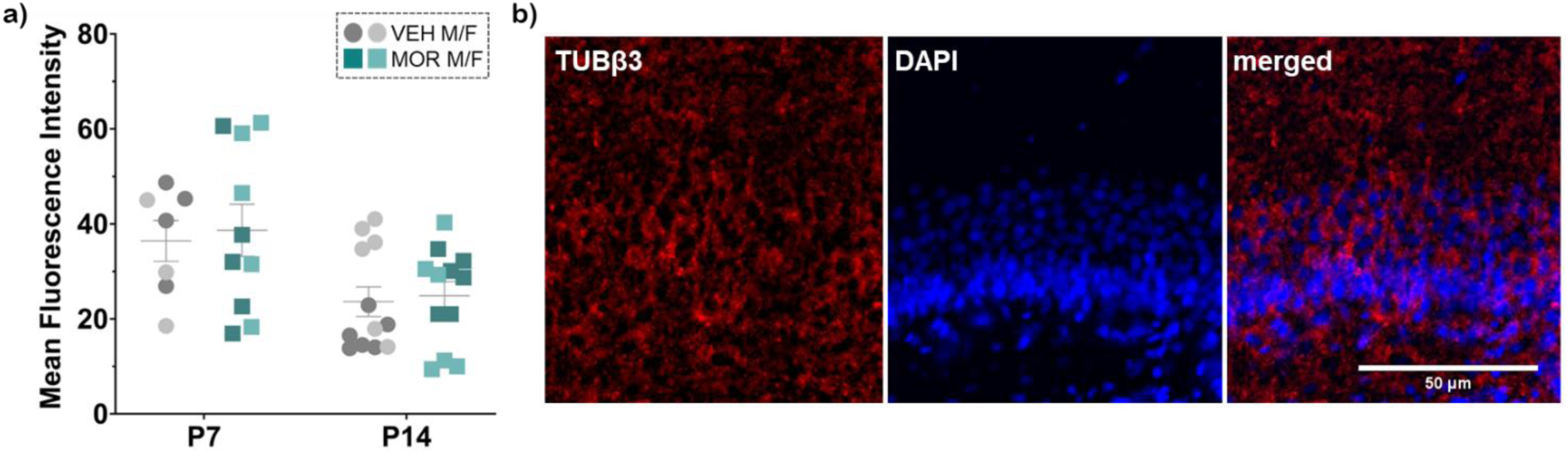
Tubβ3 expression in dorsal hippocampus (a) [n=3-6 per sex per treatment per age]; representative image of P14 DG (b)

### Neuronal Maturation

As opioids are known to direct the timing of brain development, we next assessed the effect of perigestational morphine on hippocampal neuronal maturation at multiple timepoints (P7, P14, P30, P90). Here, we used NeuN, a common marker for neurons in which high expression is limited to *mature* neurons. NeuN expression levels varied as a function of age, requiring adjustment of our image-processing protocols. Thus, we were unable to statistically compare across age.

At P7, NeuN signal within the dentate gyrus (DG) was significantly lower in MOR rats [ⴳ_VEH_=3.3% vs ⴳ_MOR_=1.6%; U=94.50, p=0.0191; **Figure 2**]. We report no effect of morphine exposure at P14 [t(21.43)=0.5480, p=0.5893]. At P30, NeuN signal area was approximately 2-fold higher in MOR rats [ⴳ_VEH_=7.3% vs ⴳ_MOR_=15.6%; t(14.42)=4.271, p=0.0007]. To determine whether this increase in NeuN expression extended into adulthood, a small cohort of VEH and MOR rats were assessed at P90. Here we observed VEH NeuN expression increased three-fold compared to P30 levels but remained below MOR, although much closer than at P30 [ⴳ_VEH_=16.4% vs ⴳ_MOR_=19.5%; t(6.429)=3.011, p=0.0218].

**Figure 2:**
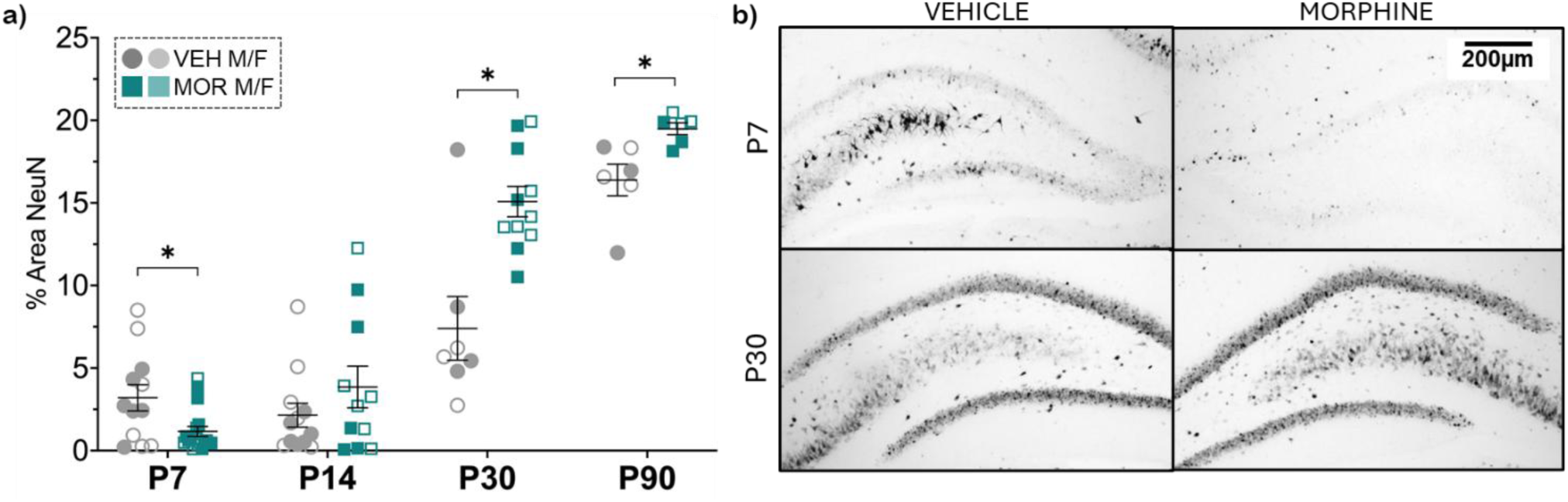
NeuN expression in DG (a) [n=3-11 per sex per treatment per age]; representative images of P7 & P30 (b); *p<0.05

### Astrocyte Proliferation

To quantify hippocampal astrocyte proliferation, we assessed co-expression of the astrocyte marker S100β and the proliferation marker Ki67 across key postnatal developmental stages (P7, P14, P30). Overall, while there was no effect of treatment on total number of S100β+ cells at P7 [F(1,10)=0.009768, p=0.9232; **Figure 3a**], the percentage of S100β+ cells that co-expressed Ki67 was significantly higher in MOR rats for both CA1 and DG regions [CA1: ⴳ_VEH_=78% vs ⴳ_MOR_=87%; DG: ⴳ_VEH_=87% vs ⴳ_MOR_=93%; F(1,10)=7.440, p=0.0213; **Figure 3b**; **Figure 4**). We also observed a significant main effect of region, with higher proliferation percentages in DG vs CA1 at this age [F(1,10)=26.30, p=0.0004].

**Figure 3:**
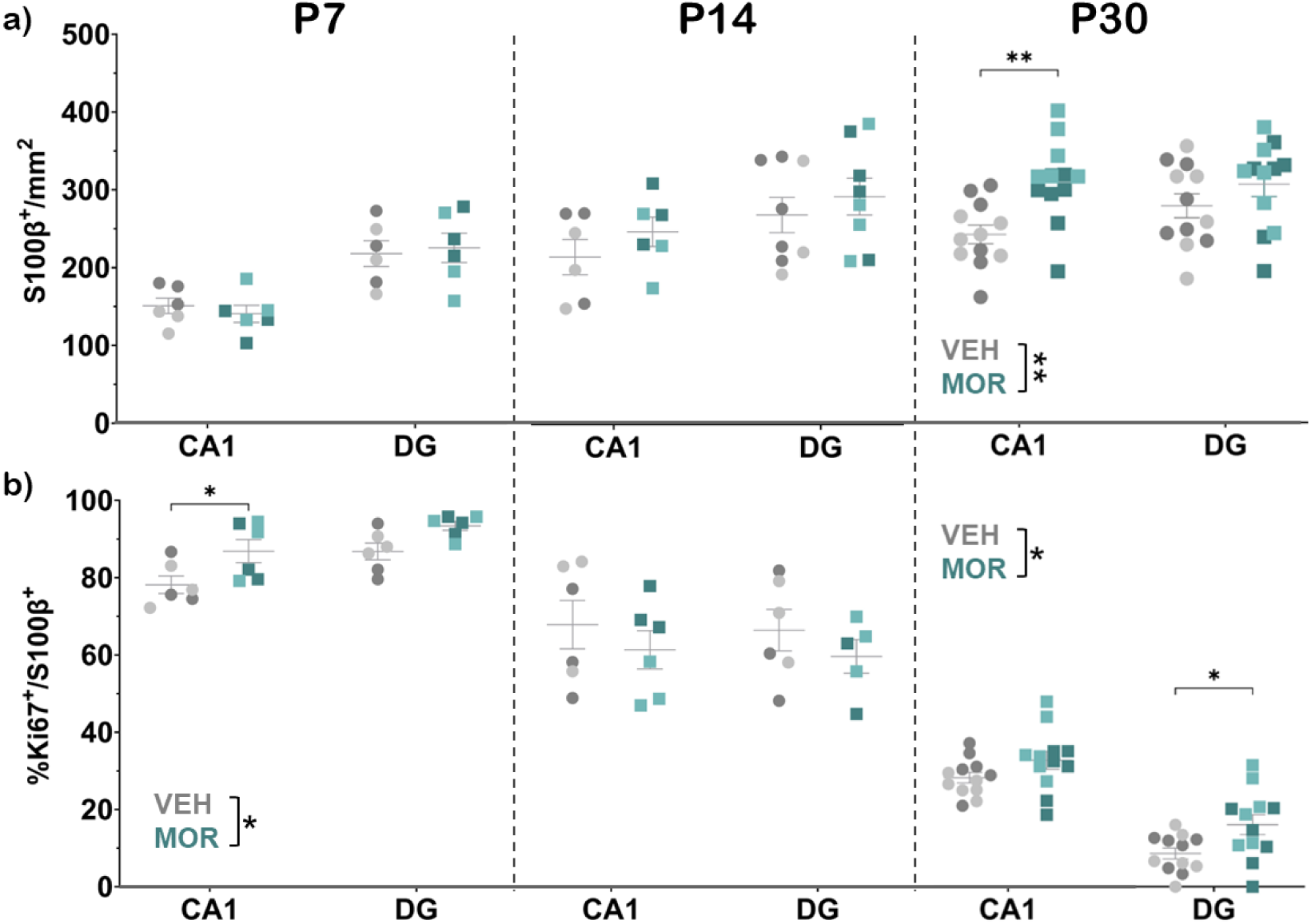
S100β expression in dorsal hippocampus at P7, P14, and P30 (a); percentage of S100β^+^ cells that co-express Ki67 at P7, P14, and P30 (b) [n=3-6 per sex per treatment per age] *p<0.05, **p<0.01

**Figure 4:**
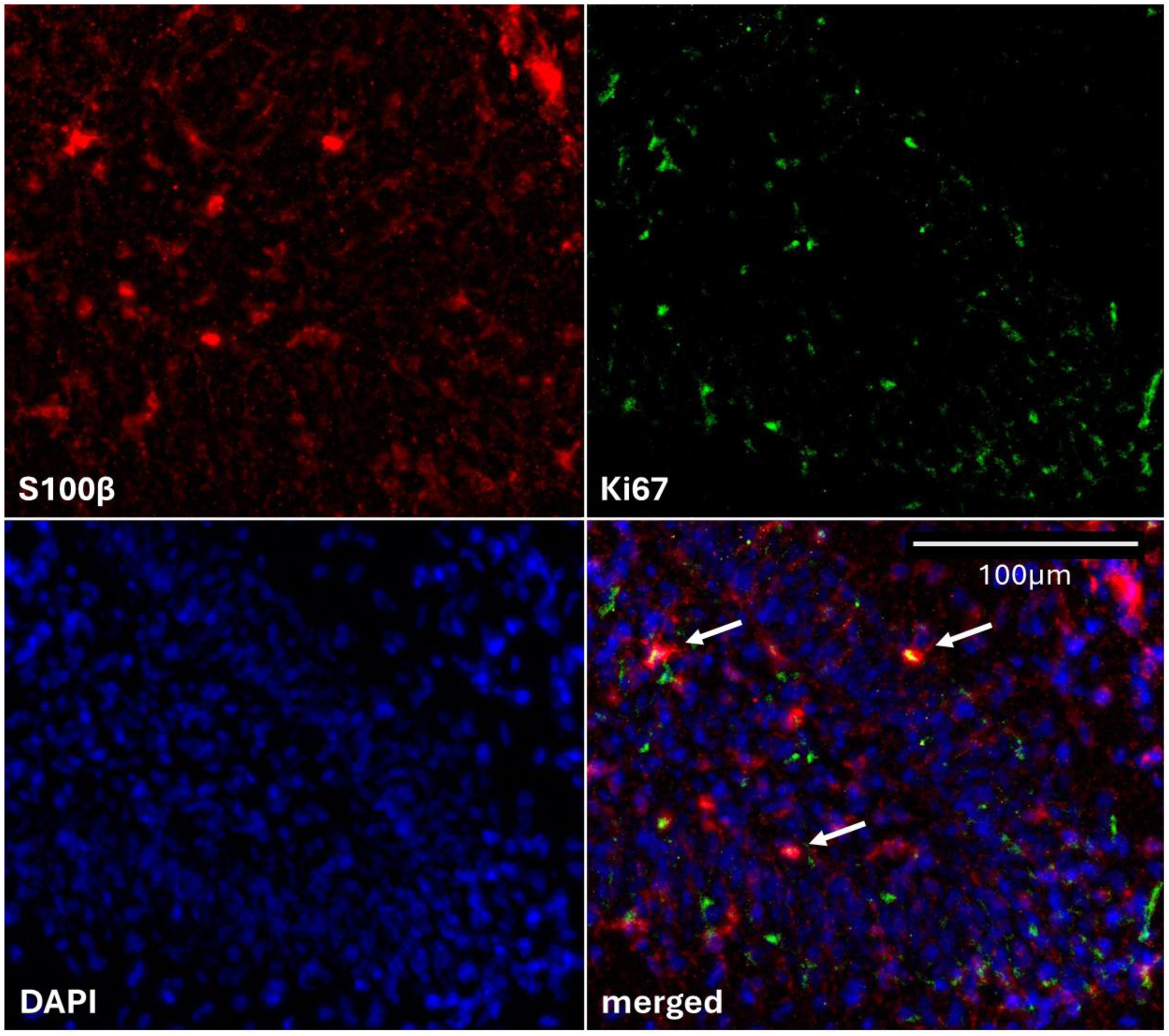
S100β/Ki67 representative images of P7 DG; white arrows indicate examples of S100β+/Ki67+ double positive cells

No effect of treatment was observed in either astrocyte count or proliferation rate at P14. [F_count_(1,24)=1.498, p=0.2329; F_proliferation_(1,10)=1.256, p=0.2886]. At P30, both astrocyte count and percent co-expression with Ki67 were significantly higher in MOR rats [Count: ⴳ_VEH_=261 vs ⴳ_MOR_=310; F_treatment_(1,22)=10.57, p=0.0037; Proliferation: ⴳ_VEH_=18% vs ⴳ_MOR_=24%; F_treatment_(1,22)=5.114, p=0.0340]. Overall, proliferation rates were significantly higher in CA1 vs DG [F(1,22)=330.6, p<0.0001].

### Oligodendrocytes

We next examined the potential impact of perigestational morphine on myelination capability, focusing on oligodendrocyte progenitor cells and differentiated oligodendrocytes that expressed Olig2. Total Olig2+ cell counts were quantified in the hippocampus at P14 and P30. Notably, we did not quantify oligodendrocytes at P7 due to the low expression of Olig2 at P7, and counts from DG and CA regions were combined as no region effects were observed. Although no treatment difference was observed at P30, cell counts at P14 were 52% higher in MOR rats compared to VEH [ⴳ_VEH_=88 vs ⴳ_MOR_=134; F_P14_(2,31)=7.199, p=0.0027; t_P30_(19.89)=0.3156, p=0.7556; **Figure 5**].

**Figure 5:**
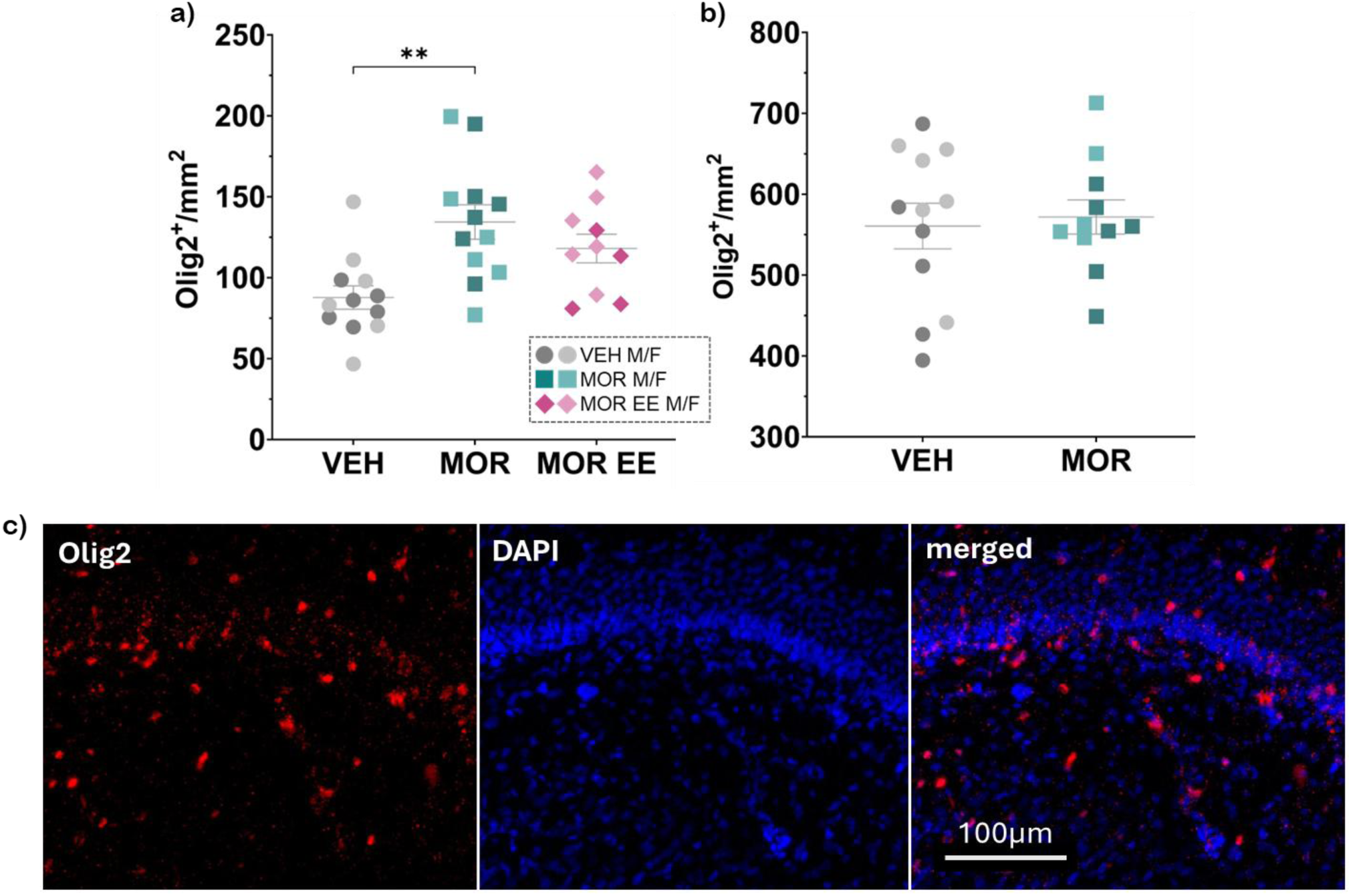
Olig2 expression in dorsal hippocampus at P14 (a) and P30 (b) [n=4-6 per sex per treatment per age]; representative images of P14 CA1 (c); **p<0.01

Clinically, infants with prenatal opioid exposure show significant alterations in white matter networks that are implicated in neurodevelopmental delays (Merhar et al., 2019; Nguyen et al., 2019; Merhar et al., 2025). Thus, we next examined if EE exposure would reverse our observed increase in Olig2 expression. Olig2+ cell counts in MOR EE rats that received home cage enrichment from P7 – P14 were 12% lower than MOR rats (ⴳ_MOR EE_=118) and not statistically different from either MOR or VEH [VEH vs MOR, p=0.0021; VEH vs MOR EE, p=0.0678; MOR vs MOR EE, p=0.4328].

### Brain-Derived Neurotrophic Factor (BDNF)

To provide a general measure of growth and development, we next examined the ratio of mature BDNF to proBDNF in the hippocampus at P14; a higher ratio denotes a more mature BNDF profile. We report a significant main effect of treatment and a significant treatment x sex interaction [F_treatment_(2,33)=6.008, p=0.0060; F_treatment x sex_(2,33)=3.425, p=0.0445; **Figure 6a**]. Perigestational morphine exposure resulted in a less mature BDNF profile in males (MOR M vs VEH M, p=0.0019) while having no significant effect in females. Notably, the change in BDNF ratio observed in males was driven by a decrease in mature BDNF [F_treatment_(2,33)=7.640, p=0.0019; F_treatment x sex_(2,33)=3.290, p=0.0498; MOR M vs VEH M, p=0.0004; **Figure 6b**]. No treatment effect was observed on proBDNF levels [F(2,33)=0.7555, p=0.4777; **Figure 6c**]. As environmental enrichment (EE) is known to boost levels of BDNF, a subset of MOR rats received home cage enrichment from P7 – P14. As shown in Figure 5a, EE successfully rescued the observed deficit in MOR males, returning their BDNF ratio to the level of controls (MOR M vs MOR EE M, p=0.0482; VEH M vs MOR EE M, p=0.6825).

**Figure 6:**
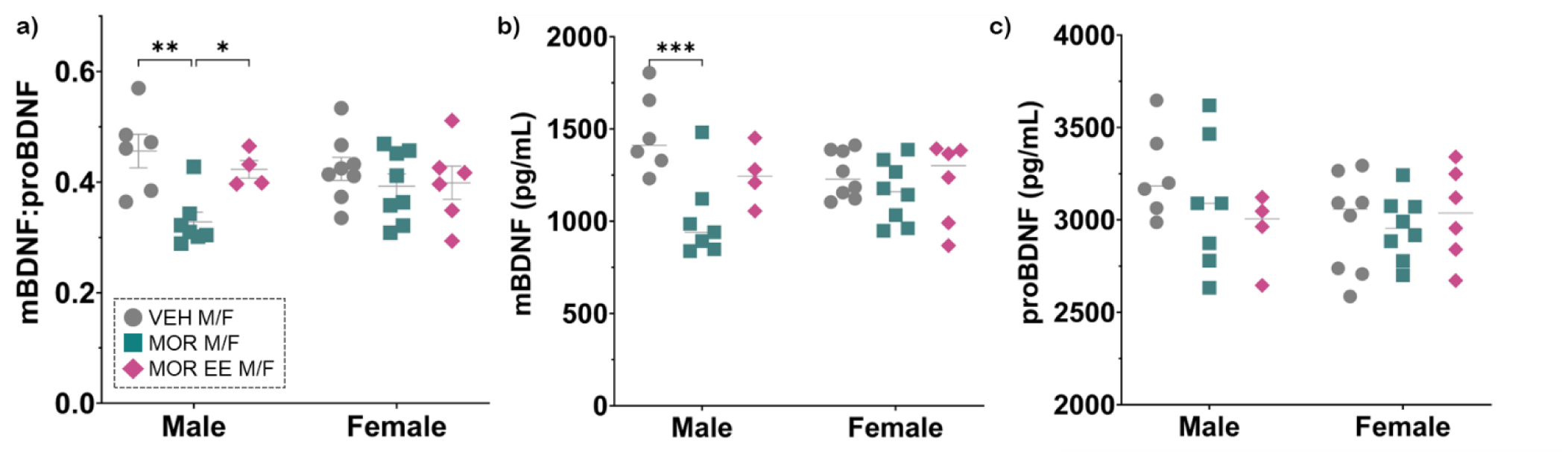
ratio of mature BDNF to proBDNF in P14 hippocampus (a); mature BDNF level (b); proBDNF level (c) [n=4-8 per sex per treatment]; *p<0.05, **p<0.01, ***p<0.001

## Discussion

The current study sought to characterize the consequences of perigestational morphine exposure on postnatal hippocampal development in male and female rats. We also examined the potential of environmental enrichment to rescue observed deficits. Here, we report that perigestational morphine exposure delayed postnatal neuronal maturation in the hippocampus with no effect on neurogenesis. We also report significant impacts of perigestational morphine on hippocampal astrocyte and oligodendrocyte proliferation, some of which were age- and region-specific. Furthermore, MOR males showed an immature hippocampal BDNF profile at P14, an effect that was reversed by home cage enrichment. In contrast, environmental enrichment had no impact on the Olig2+ cell population.

In the present study, we found no effect of perigestational morphine exposure on hippocampal neurogenesis at P7 & P14 as measured by beta-tubulin-3 (Tubβ3) expression. This lack of treatment effect was unexpected as preclinical studies have established that opioids (morphine, buprenorphine, methadone) inhibit proliferation of neural stem cells *in vitro*, and prenatal exposure to buprenorphine in rats decreases expression of neuronal markers at P21 (Wu 2014, Kibaly 2018, Boggess 2020). Furthermore, P14 MOR males in the current study displayed a significant reduction in mature BDNF, which is known to promote neural stem cell proliferation and neuronal differentiation in the hippocampus. However, while this neurogenesis-promoting effect of BDNF is well supported in adulthood, its role in neuronal proliferation during the postnatal period has garnered both supporting and refuting evidence over the years (Choi et al., 2009; Ferreira et al., 2018; Ribeiro et al. 2021). Our data suggest a limited role of hippocampal BDNF in neuronal proliferation during the postnatal period.

Although our analysis revealed no treatment effect on hippocampal neurogenesis, we did find significantly lower expression of the mature neuron marker NeuN at P7 in MOR rats. As P7 represent the end of our morphine-exposure timeline, this decrease in maturation aligns nicely with the known effect of morphine to inhibit cell growth and maturation in the hippocampus (Kibaly et al., 2019). At adult timepoints (P60 and P90), morphine-exposed males and females had significantly more NeuN than VEH, suggesting increased neuronal maturation. Alternatively, as our NeuN images were quantified as the percentage of area within the region of interest that contained signal, the observed increase in the MOR rats may indicate aberrantly located granular cells outside the granule cell layer (Amaral et al., 2007). Similar patterns of “ectopic” granule cells in the polymorphic layer of the hilus have been reported following experimentally-induced seizures and traumatic brain injury in rats, but the functional consequences of such atypical morphology is currently unknown (Scharfman 2007, Ibraham 2016). Furthermore, while preliminary counts of cells in the polymorphic layer are indeed nearly three times higher in MOR rats (7.9 VEH vs 21.6 MOR; data not shown), current analysis is unfit to identify these cells in more detail. Future investigations could further characterize these excessive neurons of the polymorphic layer, whether ectopic granule cells, excitatory mossy cells, or inhibitory interneurons, and shed more light on the potential functional significance of their increased presence following morphine exposure. This observed postnatal decrease in neuronal expression following by an increase during adolescence is of particular interest considering that, clinically, infants with prenatal opioid exposure have reduced total brain volumes and reduced gray matter volume, suggesting a decrease in neuronal density (Wu 2025).

Following investigation of neurogenesis and neuronal maturation, we assessed astrocyte and oligodendrocyte proliferation to gain a more complete picture of how postnatal hippocampal development is impacted by perigestational morphine. Astrocyte proliferation rates were significantly higher at P7 and P30, with MOR rats showing an increased proportion of cells co-expressing S100β and Ki67. P30 MOR rats also had higher S100β+ cell counts, particularly in CA1. These results indicate that morphine exposure during development may influence neural precursor cells towards glial, rather than neuronal, fates – an effect that has been demonstrated *in vitro* (Xu et al., 2015; Zhang et al., 2016). Furthermore, while astrocytes are not commonly studied in the context of prenatal opioid exposure, one recent study by Niebergall et al. (2024) reported abnormal astrocyte morphology and protein expression in cortical cultures taken from P1 rats and mice following prenatal buprenorphine exposure. Chronic opioid exposure has also been shown to reduce astrocyte-mediated synaptogenesis *in vitro*, which may underlie aberrant connectivity observed in humans following gestational opioid exposure, although no clinical reports have investigated this relationship (Ikeda 2010, Boggess 2020, Niebergall 2024). Recently, evidence has been mounting that suggests astrocytes may play a larger role in cognitive than previously believed, regulating synaptic plasticity and memory consolidation (Santello et al., 2019; Manninen et al., 2020; Escalada et al., 2024). Thus, the present report of increased astrocyte proliferation and previous reports of altered astrocyte functionality described above provide evidence of potential mechanisms underlying clinical cognitive deficits following gestational opioid exposure that warrant further investigation.

In line with the observed increases in astrocyte proliferation in the hippocampus following perigestational morphine exposure, we also report an increase in early oligodendrocytes – cells responsible for myelinating the brain – and their progenitors at P14. In line with our results, Sanchez et al. (2008) reported increased expression of myelin basic protein at P12, P19, and P26 following perigestational exposure to buprenorphine and thicker myelination within the corpus callosum at P26. Similar findings were reported in rat pups prenatally exposed to methadone, as well (Vestal-Laborde 2014). In the current study, the observed increase in Olig2+ cells did not extent into our adolescent timepoint of P30, however, Olig2 is primarily expressed by oligodendrocyte progenitor cells and recently differentiated oligodendrocytes. Future studies using our perigestational morphine exposure model will attempt to replicate reported increases in myelination during early adolescence and expand investigation to global myelination tracts in the brain using diffusion tensor imaging to further characterize the nature and extent of myeline disruption following developmental opioid exposure. A thorough preclinical characterization is of particular relevance considering that, clinically, infants with gestational opioid exposure display general white matter abnormalities and punctate white matter lesions, which have been associated with neurodevelopmental impairments (Merhar et al., 2019; Nguyen et al., 2019; Merhar et al., 2025).

Notably, one limitation of the current study is the lack of characterization of the hippocampal microglia population. Previous work from the lab has shown that perigestational opioid exposure increases microglia deramification in the periaqueductal gray following an immune challenge in adulthood, suggesting that gestational opioid exposure has a long-term impact on microglia’s reactivity and may be influencing these cells during brain development as well (Harder et al., 2025). Indeed, other forms of early-life adversity, such as limited bedding and nesting paradigms, have been shown to disrupt microglia-dependent synaptic pruning in the hypothalamus during the early postnatal period in mice (Bolton et al., 2022). Future experiments will investigate the impact of perigestational morphine to induce similar changes in microglia function during development, providing potentially yet another mechanism through which opioid exposure may be hindering cognitive performance in adolescence.

In addition to cell-specific experiments described above, we also quantified hippocampal levels of the growth factor BDNF at P14 and report a sex-specific treatment effect in which morphine-exposed males displayed an immature BDNF profile and morphine-exposed females were no different than controls. Our results are similar to those of Wu et al. (2014) who reported reduced whole-brain BDNF expression in both male and female rats at P21 following gestational buprenorphine exposure as well as decreased phosphorylation of the BDNF receptor TrkB. Reductions in hippocampal BDNF were also reported in late adolescent rats (P51) following prenatal morphine exposure (Ahmadalipour 2018). One notable difference between the current and previous reports is the quantification of both mature BDNF and its active precursor, proBDNF, in the present study, which allows for a more nuanced BDNF ratio rather than raw BDNF expression. Higher levels of proBDNF to mature BDNF in the hippocampus can negatively impact synaptic plasticity and alter cognitive processes mediated by the region, such as spatial learning and memory consolidation, highlighting the need to quantify both (Yang et al., 2015).

The current report of decreased BDNF maturity in MOR males aligns nicely with recently published data from the lab showing MOR rats have impairments in spatial learning, preferring a serial strategy over a more efficient, hippocampus-dependent spatial strategy in the Barnes maze (Vogt & Murphy, 2025). Deficits in both measures were rescued in males by environmental enrichment, suggesting in agreement with other rodent studies that increases in BDNF can improve performance on spatial learning tasks due to its role in regulating activity-driven synaptic plasticity (Mizuno et al., 2000; Colucci-D’Amato et al., 2020). Notably, however, the previously reported impairment in spatial learning was displayed by MOR males and females while only MOR males show BDNF deficits in the current study. Underlying sex differences in hippocampal BDNF expression, BDNF-TrkB signaling, and behavioral outcomes of BDNF knockouts have been established in several rodent models, likely driven by sex-steroid-mediated regulation of BDNF signaling, as reviewed by Chan and Ye (2017). Thus, we hypothesize that the molecular basis for spatial learning in females may be less dependent on BDNF than it is in males. Future experiments including mechanistic manipulations will further elucidate the differences in neurocircuitry that underly these sex-specific results.

In conclusion, our findings indicate that perigestational morphine exposure delays postnatal neuronal maturation and increases astrocyte and oligodendrocyte proliferation in the hippocampus. We further report an immature BDNF profile in morphine-exposed males but not females that is rescued by home cage enrichment. These results underscore developmental consequences of opioid use during pregnancy and highlight environmental enrichment as a promising non-pharmacological approach to support brain growth recovery in offspring exposed to opioids *in utero*.

## Acknowledgements

This work was supported by the National Institutes of Health (AZM; R34DA061483), Georgia State University’s Brains and Behavior Program (MEV), and the Beckman Scholars Program (JK). The funding sources were not involved in study design, data collection, analysis or interpretation, manuscript writing, or the decision to publish this article. We gratefully acknowledge the NIDA Drug Supply Program for providing morphine sulfate.

## Notes

### Competing Interest Statement

The authors have declared no competing interest.

